# Clustering Gene Co-expression Using 2D Contour Analysis and Autoencoder-based Embedding

**DOI:** 10.1101/2025.09.07.670504

**Authors:** Vikraman Karunanidhi, Ken-ichi Aisaki, Jun Kanno, Natalia Polouliakh, Samik Ghosh, Hiroaki Kitano, Sucheendra K Palaniappan

## Abstract

Identifying co-expressed genes is crucial for understanding biological processes; however, common methods such as Pearson and Spearman correlation rely on assumptions of linearity and monotonicity, respectively, that may not hold for complex biological data. To address these limitations, we propose a framework that captures visual features of gene expression profiles without relying solely on correlation-based methods. Our approach involves converting 3D gene expression data, which contain rich information on gene envelopes over time, into 2D contours that retain important visual information. We then train an autoencoder on the 2D contour images and generate embeddings from them, followed by clustering on the generated embeddings. We also introduce a new clustering algorithm, TAHC, based on hierarchical clustering, which performs better than existing methods in higher dimensions while using cosine similarity. We apply this framework to the Percellome database, which contains gene expression data with two variables—time and dosage—across various experimental conditions (e.g., tissue type). The resulting clusters exhibit good visual coherence, with an overall average Pearson correlation coefficient of 0.81, demonstrating the effectiveness of our approach.

## 1 Introduction

Identifying co-expressed genes is a critical step in understanding biological processes, as such genes are likely to be part of related pathways and regulated by similar mechanisms (1). Discovering co-expressed genes forms the basis for constructing gene co-expression networks, which are invaluable for associating genes with specific biological functions, pathways, and other important cellular activities.

In this paper, we work with the Percellome project(2), which is a comprehensive database of gene expression profiles from experiments carried out under various chemicals, tissues, organs, and experimental conditions in three different groups. For each experiment, gene expression data are measured across multiple dosage levels and time points. In the Percellome project it is conceptualized that a set of co-expressed genes activated at a specific time point can trigger another set of co-expressed genes at a later time point, resulting in a ripple effect (2). With this extensive database, comparing co-expressed gene sets across different chemicals and biological conditions offers the potential to uncover critical insights into underlying biological processes.

The primary objective of this paper is to reliably identify co-expressed genes from a single experiment of Percellome database,laying the groundwork for scaling the analysis to comparisons across multiple experiments.

Currently, analysis of the Percellome database often involves manual inspection of 3D surface plots of gene expression profiles. However, with each experiment encompassing up to 45,101 gene expression values measured across multiple dosage levels and time points, and a total of approximately 370 experiments spanning various tissues and chemicals, this manual approach is laborious. Therefore, an automated system is required to capture the visual characteristics of gene expression profiles and group genes.

Genes that have similar gene expression profiles are likely to be co-expressed. Therefore, to find co-expressed genes, we need to find genes that share similar gene expression profiles. While many existing methods are available for identifying co-expressed genes, most focus solely on raw amplitude values and fail to consider visual features present in the gene expression plot. Commonly used approaches for gene co-expression analysis (3, 4) rely on correlation methods such as Pearson correlation, which may fail to capture complex patterns that are biologically significant.

To address these limitations, we propose a new approach that captures the visual features of gene expression profiles represented in 3D surface plots and clusters them without relying on correlation-based methods. To extract these visual features, we convert 3D gene expression profiles into 2D contour plots, which are subsequently transformed into embeddings using sparse autoencoders. Variability is incorporated by calculating the standard deviation at peak expression values across triplicates, which is then combined with the embeddings for clustering. Using these combined features, we cluster genes into groups that share similar expression profiles and comparable variability ranges.

Beyond this immediate application, our long-term objective is to investigate how gene expression patterns vary across tissues, chemical treatments, and experimental conditions. For this purpose, cosine similarity is particularly effective, as it quantifies the similarity of gene expression profiles both within and across experiments. As part of this work, we developed a clustering algorithm optimized for high-dimensional embeddings using cosine similarity.

The main contributions of this paper are as follows:

- A representation of 3D gene expression profiles that captures visual features using autoencoders and 2D contour plots.
- A clustering approach based on hierarchical clustering that shows improved performance over other algorithms for clustering embeddings of 2D contours, using the cosine similarity metric.

We begin by reviewing related work on gene co-expression analysis, the application of autoencoders in biological contexts, and clustering methodologies. Next, we provide a detailed description of the methods employed in this study. In the results section, we first determine the optimal parameters for our autoencoder and select the best-performing model for clustering. We then evaluate the performance of different clustering algorithms within our use case. Finally, we analyze the clusters generated by our approach using one of the experiments from the Percellome project.

## 2 Related work

Gene expression plots capture rich information about the landscape of how genes respond to different conditions like dosage and time. Commonly used methods to find co-expressed genes based on gene expression values rely on correlation-based methods like Pearson correlation and Spearman correlation. However, Pearson Correlation expects normality in data and might not be suitable for biological data and Spearman correlation expects monotonic relationships (3),(5). In some cases Pearson Correlation may still be preferred over Spearman correlation (5).

To address these statistical constraints, an alternative is to use representation learning techniques that can extract richer, multidimensional features from gene expression data. Autoencoders, in particular, are well suited for capturing complex patterns (6) because they learn data-driven representations that preserve non-linear relationships, going beyond what is possible with simple correlation measures.

Autoencoders(7) are a type of unsupervised neural network that learn compact, efficient representations of data by encoding high-dimensional inputs into a lower-dimensional latent space and reconstructing them back. The two main components are an encoder that transforms the input into a reduced representation, and a decoder that reconstructs the original input from this representation. The latent space often preserves meaningful structure in the data, which can later be used for downstream tasks (8). Autoencoders have many applications (9), including dimensionality reduction, feature representation, anomaly detection, and image denoising and can be applied to various data modalities, including omics data(10), images, text and time series (11).

Besides reconstruction, a variant known as the variational autoencoder (VAE) (12) learns a probabilistic latent space, enabling the generation of new data samples by decoding from the learned distribution.

Autoencoders offer major advantages in biological contexts, enabling the condensation of diverse biological information—such as gene expression profiles, and multi-omics signals—into compact latent representation (13–15).

P. Wickramasinghe et al (16) used to autoencoders to reduce dimensions of gene sequence data. Wang, D., & Gu (17) used variational autoencoders to visualise Single-cell RNA sequencing (scRNA-seq) datasets and applied their technique to identify several candidate marker genes associated with early embryo development. Zhang, et al(18) used graph based autoencoders to integrate gene expression, cellular neighborhoods, and chromatin imaging in a joint latent representation and used it for studying the spatio-temporal progression of Alzheimer’s disease.

Many variants of autoencoders have been proposed since its inception. Sparse Autoencoders (19),which we employ in this work, learn efficient and meaningful representations by enforcing sparsity in the latent representation. Recently, they have been extensively used for interpretability of LLMs (20, 21). They do so by using overcomplete autoencoders that have more number of neurons in the hidden layer than input dimension and using sparse loss function. By inducing sparsity and having a large number of neurons in the hidden layer, they find which neurons activate for a particular concept, thereby finding interpretable features.

A representation of gene expression data enables the comparison of changes in expression levels across experiments under varying conditions such as tissue type, chemical treatment, and other factors. Additional experimental metadata can also be incorporated into the latent representation for more comprehensive analysis. In this work, we generate such representations using sparse undercomplete autoencoders trained on 2D contour plots of 3D gene expression profiles, and we incorporate gene variability information alongside the embeddings for clustering genes that have similar expression profiles.

Agglomerative clustering, a form of hierarchical clustering, is commonly used in gene expression analysis. However, in our experiments, the clustering of embeddings with a fixed threshold and cosine similarity led to the merging of distinct clusters. To address this, we developed Threshold Adaptive Hierarchical Clustering (TAHC), a variant of agglomerative clustering that adaptively updates the threshold at each iteration and halts merging once the criteria are met. TAHC shares a conceptual similarity with the adaptive mean-linkage algorithm (22), which also recalculates merging thresholds based on local distance statistics.

## 3 Methods

### 3.1. Dataset

In this section, we provide an overview of the dataset and the techniques used in its processing and generation. For training our autoencoder model to generate latent representations of the 2D contours, we used gene expression data from two experiments in the Percellome Database. For training and validation, we used data from the TTG020-L experiment, and data from the TTG016-L(C) experiment (23) for testing. Gene expression values were collected under different dosage levels and time intervals. In the TTG020-L experiment, data were collected from five dosage values: 0, 1.23, 3.70, 11.11, and 33.33 µg/kg, which were converted into categorical labels (V, L, ML, MH, and H) corresponding to different intensity levels. In the TTG016-L(C) experiment, data were collected from four dosage values: 0, 10, 30, and 100 units, which were also assigned labels (V, L, M, and H). In both datasets, gene expression was measured at 2, 4, 8, and 24-hour intervals, with the time points also converted into categories for plotting purposes.

#### 3.1.1. Data representation using 2D contours

2D contour plots (Fig.1) are a two-dimensional representation of three-dimensional data, where the third dimension is encoded as intensity or color values in the image. These plots effectively capture the overall shape, peaks, gradients, and other critical features present in the original 3D data, enabling a comprehensive visualization of complex relationships. This representation is particularly advantageous for use with 2D convolutional neural networks, as it retains the spatial structure and visual artifacts of the data. By preserving key visual features, these plots facilitate the extraction of biologically meaningful patterns and enable downstream clustering and analysis of co-expressed genes.

**Fig. 1.**
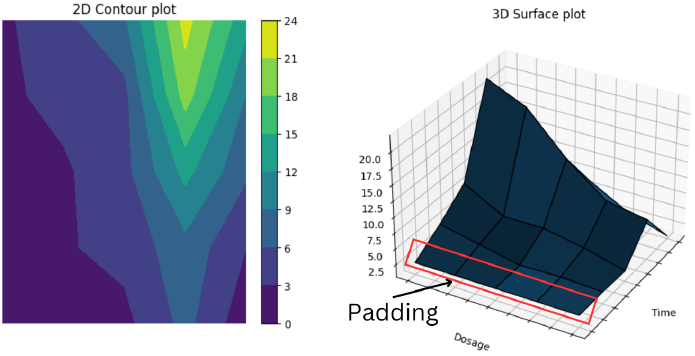
A sample 2D contour plot and corresponding 3D surface plot

#### 3.1.2. Data processing and Generation

To prepare the gene expression data for downstream tasks such as representation learning and clustering, we apply a series of preprocessing steps, including padding, averaging, and contour plot generation. These steps aim to standardize the data format and enhance interpretability of temporal and dosage-related gene expression patterns.

##### Padding

To improve the visualization of changes at the 2-hour mark, we pad the value at 2 hours and zero dosage across all dosage points as a reference (23). The padded value acts as a virtual zero-hour baseline. After applying the virtual zero-hour padding, the probe expression data in the TTG020-L experiment, which includes 5 dosages and 4 time points, is transformed into a 5×5 matrix, whereas the data in TTG016-L(C), which contains 4 dosages and 4 time points, is transformed into a 5×4 matrix.

##### Steps involved in generating 2D contours

- Calculate the mean gene expression value across triplicates for each time-dosage point.
- Pad values with virtual zero-hour values.
- Generate 2D contour plots using the processed values.

##### 2D Contour Plot Configuration

- The X and Y axes correspond to time and dosage categories, respectively, and are equally spaced. Equispaced categorical values are used to remain consistent with the methodology in (2).
- Color is encoded to reflect the intensity of gene expression at each time-dosage point.
- A consistent color map is applied across all plots.

#### 3.1.3. Synthetic data generation

To evaluate the quality of the latent representations produced by our model under different hyperparameters, we created a synthetic dataset consisting of 2D contour plot images and their corresponding cluster labels. This synthetic dataset also enabled the evaluation and comparison of the performance of different clustering approaches.

We selected 16 distinct gene expression profiles, ensuring they were sufficiently varied from one another. To augment the data, we added Gaussian noise (Eq. (1)) to the average gene expression values for each of the 16 profiles. We limited the mean and standard deviation of the noise to small values (*µ* ∈ [−0.2, 0.2], *σ* ∈ [0.16, 0.20]) to ensure that the original signal was not significantly altered. These values were chosen to maintain the identity of each cluster in the synthetic dataset, preventing overlap between distinct clusters and ensuring the augmented samples remained close to their source profiles. Consequently, the embedding vectors for data within the same group should exhibit high similarity and should be closer together in latent space. In total, we generated 3,200 image-label pairs, comprising 200 samples for each of the 16 selected gene profiles, resulting in 16 unique labels.

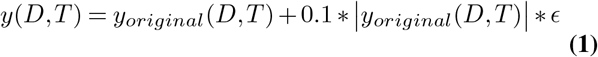

where:

- *y*_*original*_(*D, T*) is the original signal at Dosage = D and Time = T,
- is the noise term, sampled from a Gaussian distribution *N* (*µ, σ*^2^),

### 3.2. Model architecture and training details

To effectively capture the features of 2D contour plots, we used sparse autoencoders (SAEs) to learn rich latent representations. We chose SAEs over standard autoencoders because they enforce sparsity in the latent space, encouraging the model to learn features from a limited number of active neurons. This is achieved by incorporating an additional regularization term in the loss function that penalizes activations in the latent layer.

The SAE used in this work, like a standard autoencoder, comprises an encoder and a decoder. Since the input is an image, we used convolutional layers in the model architecture to effectively capture spatial features. The encoder comprises convolutional layers with a stride of 2 and a kernel size of 3, each followed by an ELU activation function. We restricted the encoder depth to 3 and 5 convolutional layers, as the first few layers in CNNs are known to capture the most essential shape-related features such as edges and contours (24). In the three-layer configuration, the number of channels is set to 32, 64, and 128, while in the five-layer configuration, the channels are set to 32, 64, 128, 256, and 512. This setup follows common CNN design conventions, where the number of channels increases with depth (25). At the end of the convolutional layers, the outputs are flattened and passed through a linear layer, which downsizes them into the bottleneck layer to form the latent representation.

The decoder has a linear layer that upsamples the latent representation from the encoder. The output of this linear layer is reshaped to match the pre-flattening dimensions of the final convolutional layer in the encoder. Following this, the decoder employs upsampling layers with the number of channels mirroring the encoder: 128, 64, 32 for the three-layer configuration and 512, 256, 128, 64, 32 for the five-layer configuration. The stride, kernel size, and activation function in the decoder are identical to those in the encoder. The input to the model has a shape of 224 × 224 × 3, with pixel values normalized to fall within the range of 0 to 1. The sigmoid activation function was used for the final layer to ensure that the output values are restricted to the range of 0 to 1, consistent with the input range. The model was trained for 100 epochs using the Adam optimizer with a learning rate of 1 × 10^−4^, and a learning rate scheduler that reduced the learning rate by a factor of 0.2 if the validation loss did not improve for 5 consecutive epochs, with a minimum learning rate of 5 × 10^−5^.

#### 3.2.1. Loss functions

Since the proposed model is a sparse autoencoder, the loss function includes both reconstruction error and a sparsity regularization term, and is defined as follows:

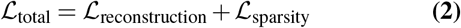

Where:

1. Reconstruction loss:

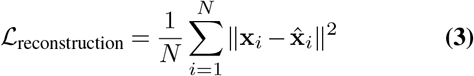 Here, **x**_*i*_ is the original *i*-th input image, 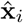 is the reconstruction of the input image, and *N* is the number of samples. The reconstruction loss is the Mean Squared Error between the original image and the reconstructed image.
2. Sparsity loss:

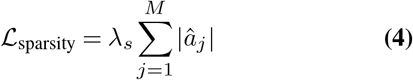

where 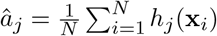 is the average activation of the *j*-th hidden unit in the latent layer, and *λ*_*s*_ is the regularization constant (sparsity coefficient) controlling the sparsity strength. combining Eq. (3) and Eq. (4), we obtain the total loss function:

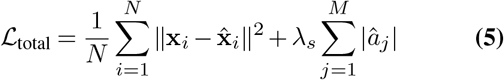

#### 3.2.2. Hyperparameters

Our primary objective in training the sparse autoencoder is to generate high-quality latent representations of 2D contours, capturing shape, curvature, and other visual features. The quality of these latent representations is primarily influenced by the following factors: the compression factor (i.e., the ratio of neurons in the latent layer to those in the flattened output of the final convolutional layer), the number of neurons in the latent layer, and the sparsity coefficient used in the loss function.

To study the effect of compression on the quality of latent representations, we varied both the number of neurons in the flattened layer (i.e., the output of the final convolutional layer) and the latent layer (i.e., the hidden layer of the autoencoder). The number of neurons in the flattened layer was adjusted by using two different convolutional neural network (CNN) configurations: one with three layers and another with five layers, while keeping other parameters such as stride and padding constant.

For each CNN configuration, we experimented with different numbers of neurons in the latent layer. In the three-layer CNN, we tested latent sizes ranging from 32 to 256 neurons, and in the five-layer CNN, we extended this range up to 512 neurons. This setup allowed us to analyze how varying compression levels—resulting from different combinations of flattened and latent layer sizes—affect the learned representations.

The sparsity regularization term in the loss function Eq. (4) encourages the model to learn sparse latent representations by minimizing hidden layer activations. To evaluate its impact, we conducted experiments using five different sparsity coefficients, including zero, and analyzed their influence on both input reconstruction quality and learned latent representations. The inclusion of a zero sparsity coefficient serves as a baseline, allowing for isolation and analysis of the effect of sparsity under different training conditions.

In summary, we vary the following parameters to evaluate the performance of our model:

- Number of neurons in the latent layer (32,64,128,256,512)
- Number of convolutional layers (3 layer and 5 layer CNNs)
- Sparsity coeeficient *λ*_*s*_ (0, 10^−2^, 10^−1^, 1, 10)

### 3.3. Clustering

Clustering is a key component of our pipeline for identifying co-expressed genes. During our initial experiments with the TTG020-L dataset, we applied dimensionality reduction to the embedding vectors and then performed clustering. However, this caused the data to become tightly clustered in the low-dimensional space, reducing inter-point distance variability. As a result, clustering algorithms like DBSCAN tended to merge most points into a single cluster and became overly sensitive to hyperparameters such as `eps` and min_samples. Based on these observations, we decided to exclude dimensionality reduction from further analysis.

Without dimensionality reduction, we experimented with both DBSCAN and agglomerative clustering using cosine similarity as the distance metric. Cosine similarity is particularly well-suited for high-dimensional embeddings because it is scale-invariant and facilitates comparison across different experimental conditions. However, DBSCAN did not perform well in this setting, as it failed to separate meaningful clusters in the synthetic dataset. Similarly, applying a fixed threshold in hierarchical clustering often resulted in suboptimal clusters in the synthetic dataset, and adjusting the threshold across runs did not consistently yield ideal groupings.

To overcome these challenges, we developed a new method: Threshold-Adaptive Hierarchical Clustering (TAHC), which dynamically adjusts the similarity threshold during the clustering process. Our adaptive method consistently produced clusters closely aligned with the ideal clusters, outperforming other methods. (sec: 4.3.1).

#### Algorithm 1

Clustering Algorithm - TAHC

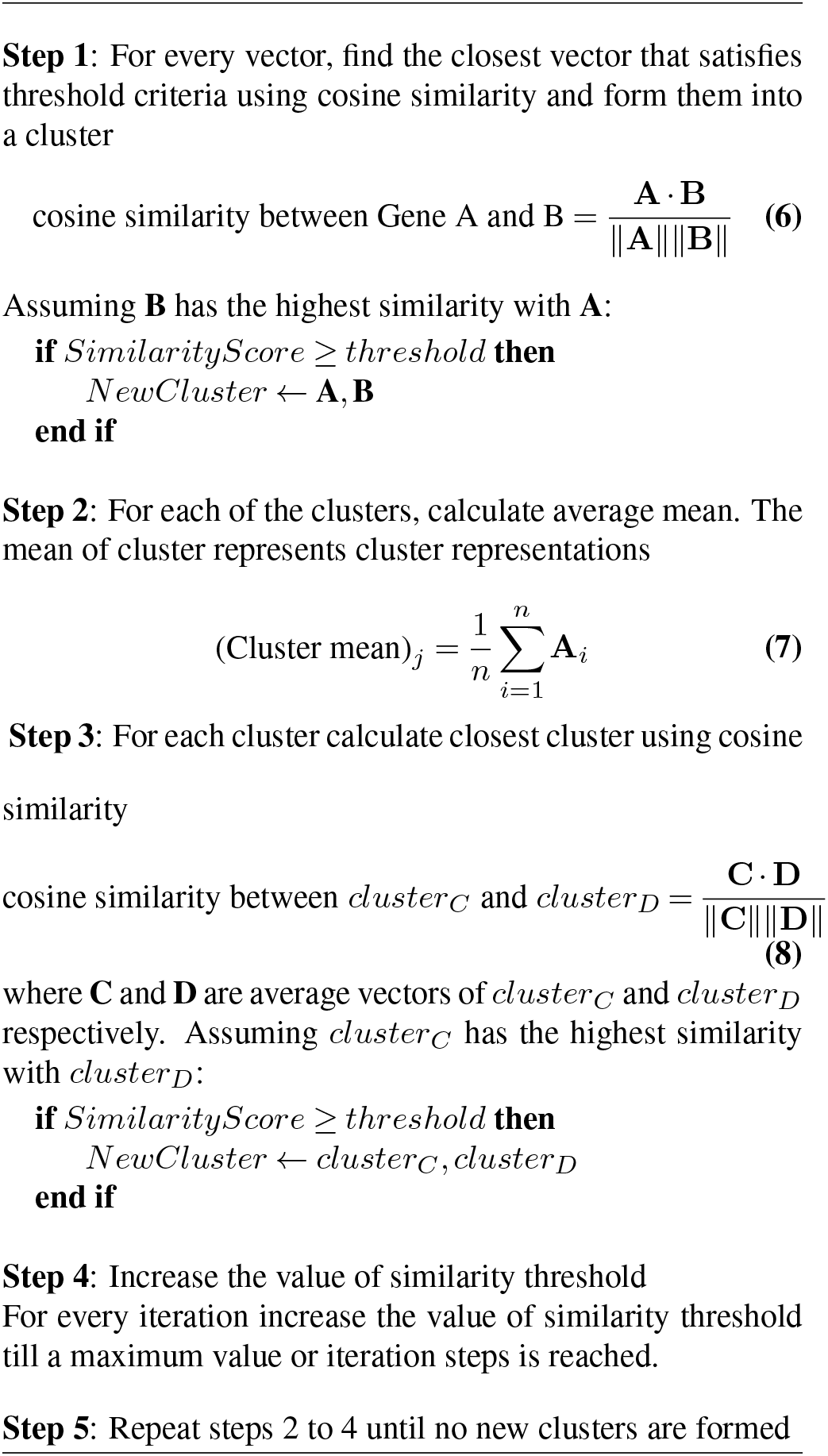

#### 3.3.1. Threshold Adaptive Hierarchical Clustering (TAHC)

Threshold-Adaptive Hierarchical Clustering (TAHC) is a variant of hierarchical clustering that introduces a dynamic thresholding mechanism to guide the merging process. Unlike traditional methods like agglomerative clustering, which typically rely on a fixed threshold to determine cluster boundaries, TAHC adaptively increases the threshold at each iteration, progressively tightening the merge condition over time. This adjustment helps mitigate unintended cluster overlap caused by averaging effects during merging, particularly in high-dimensional spaces. While conceptually related to the adaptive mean-linkage algorithm (22), which dynamically adjusts merging thresholds based on local distance statistics, TAHC differs in its formulation. In contrast to the adaptive mean-linkage algorithm’s minimax-driven, non-monotonic threshold updates over Euclidean distances, TAHC employs a progressively increasing cosine similarity threshold between cluster centroids. This threshold is increased up to a predefined limit, providing a more interpretable and tunable approach to controlling the hierarchical merge process.

To apply TAHC within our framework, we begin by extracting all embedding vectors from the encoder, which serve as input to the clustering algorithm. In the first iteration, each embedding vector is compared against all others to identify the most similar vector. If the cosine similarity exceeds the threshold, the corresponding pair is merged into a cluster. Once all possible clusters are formed, the threshold is increased.

From the second iteration onward, each cluster is represented by its mean vector, computed as the average of all embedding vectors within the cluster. For each cluster, the nearest neighbor for each cluster is computed based on mean vector similarity. If two clusters meet the updated threshold criterion, they are merged. This process is repeated for all clusters in the iteration.

After each iteration is complete, the similarity threshold is increased until either a predefined maximum value is reached, or a specified number of threshold-increasing steps have been completed. The entire process is repeated until no new clusters are formed.

### 3.4. Incorporating variability

In our dataset, variability arises from the triplicates. There are various ways to define the variability among the triplicates, and in this study, we focus on the variability at the highest value. To quantify this variability, we calculate the standard deviation across the triplicates. First, we normalize the data using Z-score normalization, which transforms the values to a mean of zero and a standard deviation of one. Normalization is essential because the maximum value range of gene expression values is widespread. By adding the absolute values of the Z scores, we obtain a single quantitative measure of overall variability among the triplicates for that particular gene. We then compute the sum of the absolute value of Z-scores which provides an overall measure of variability across the triplicates. Finally, we categorize the sum into four levels of variability: low variance, low-medium variance, medium-high variance, and high variance.

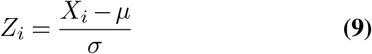

where:

- *Z*_*i*_ is the Z-score of the *i*^*th*^ group,
- *X*_*i*_ is the value from the *i*^*th*^ group,
- *µ* is the mean at peak position,
- *σ* is the standard deviation at peak position.

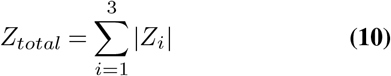

Using the values from Eq. (10), we plotted a histogram and categorized the resulting *Z*_*total*_ scores into four discrete bins. Subsequently, we converted the continuous *Z*_*total*_ values into categorical labels based on these bins, which we integrate into our pipeline.

#### Integrating standard deviation categories

Our goal is to cluster gene expression values based on both visual similarity and variability. Directly fusing the standard deviation category into the high-dimensional vector did not add much information. The cosine similarity between the same embedding with two different standard deviation categories resulted in very high similarity scores. Hence, instead of fusing the standard deviation directly, we first split the data based on standard deviation category. After splitting, clustering is performed separately for each category. In this way, clustering considers both visual similarity and levels of variability.

### 3.5. Functional Enrichment Analysis

To assess the biological relevance of the gene clusters derived from expression data, we performed functional enrichment analysis using `GSEApy` (26). This package implements the Gene Set Enrichment Analysis (GSEA) framework (27). Pathway gene sets were obtained from the `KEGG_2019_Mouse` geneset from the KEGG database (28), as the Percellome data is based on mouse experiments. Statistical significance was determined using an adjusted p-value threshold of 0.05, which controls for multiple hypothesis testing by limiting the expected proportion of false positives among the significant results. Only clusters with more than 10 genes were considered for enrichment.

### 3.6. Pipeline

Our proposed workflow consists of two primary phases: a training phase to learn compact representations of gene-expression data and an inference phase to cluster similar gene-expression profiles. The detailed steps are as follows:

- **Training phase**
  1. Extract gene expression values from the dataset and compute the mean expression value from triplicates for each dosage–time point.
  2. Generate the corresponding 2D contour images for each expression profile.
  3. Train an autoencoder using the 2D contour images to learn low-dimensional representations of features present in the plots.
- **Inference phase**:
  1. For the dataset of interest, extract gene-expression values and generate the corresponding 2D contour images.
  2. Input these images into the trained autoencoder to obtain embeddings from the encoder.
  3. Group the embeddings according to their associated standard deviation categories to incorporate variability information.
  4. Perform clustering analysis separately within each standard deviation category to identify similar gene co-expression profiles.

## 4 Results

We trained our model on an NVIDIA GeForce RTX 2070 GPU with 8GB RAM, running on an Intel Core i7 10th Gen system with 32GB RAM. Each training experiment took approximately three hours for 100 epochs. For clustering 3,200 embedding vectors of size 512, TAHC took around 3.5 seconds, while Agglomerative Clustering took about 1.5 seconds on our system.

The following steps outline the structure of our results and analysis:

1. Analyze the effect of different hyperparameters on model performance.
2. Select the optimal hyperparameters based on performance on the image reconstruction, quality of latent representation, and use the corresponding model for further analysis.
3. Compare the performance of various clustering algorithms on embeddings generated by the selected model on the synthetic dataset.
4. Apply clustering analysis on the TTG020-L experimental dataset using the selected model.

### 4.1. Effect of hyperparameters on image reconstruction

To examine the effect of latent dimension size on image reconstruction, we first report metrics without sparsity. Subsequently, we evaluate how both the sparsity coefficient and the latent dimension size influence model performance. We use Eq. (3) to calculate the reconstruction loss.

#### 4.1.1. Effect of latent dimension over image reconstruction - without sparse loss

The number of neurons in the latent dimension is crucial for generating embeddings, with each individual neuron or group of neurons likely capturing specific features of the 2D contours. To investigate the effect of the latent dimension, we set the sparsity coefficient to zero during training to ensure that sparsity loss did not influence the analysis. For both the three- and five-layer CNN models, increasing the latent dimension improved image reconstruction quality up to a certain threshold (Fig.2). For the three-layer CNN network, image reconstruction quality did not improve significantly beyond 128 dimensions (Fig.2a). Similarly, for the five-layer CNN network, no significant enhancement in reconstruction quality was observed after 256 dimensions (Fig.2b).

**Fig. 2.**
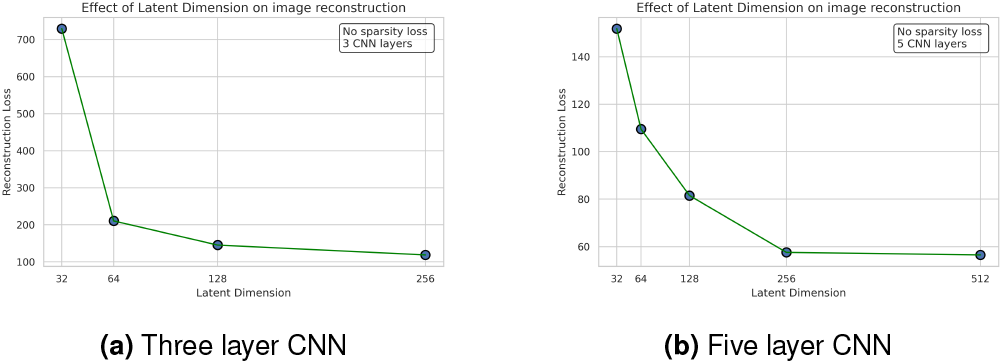
Effect of latent dimension on image reconstruction

#### 4.1.2. Effect of sparsity on image reconstruction

To study the effect of sparsity on image reconstruction, we varied the sparsity coefficient chosen from Sec. 3.2.2 across different latent dimensions. The magnitude of the sparsity coefficient affects the overall loss (Eq. (5)), with higher values imposing a stronger penalty on the activations, thereby encouraging sparsity and impacting overall reconstruction performance. In the case of the three-layer CNN, the sparsity loss consistently improves reconstruction across all sparsity coefficients, although its impact diminishes as the latent dimension increases (Fig.3a). For the five-layer CNN, an increase in the sparsity coefficient generally leads to a negative effect on image reconstruction across various latent dimensions. However, for the latent dimension of 512 (Fig.3b), all sparsity coefficients—except for the coefficient of 10—seem to converge to a similar value, which corresponds to the lowest reconstruction loss (Fig.3).

**Fig. 3.**
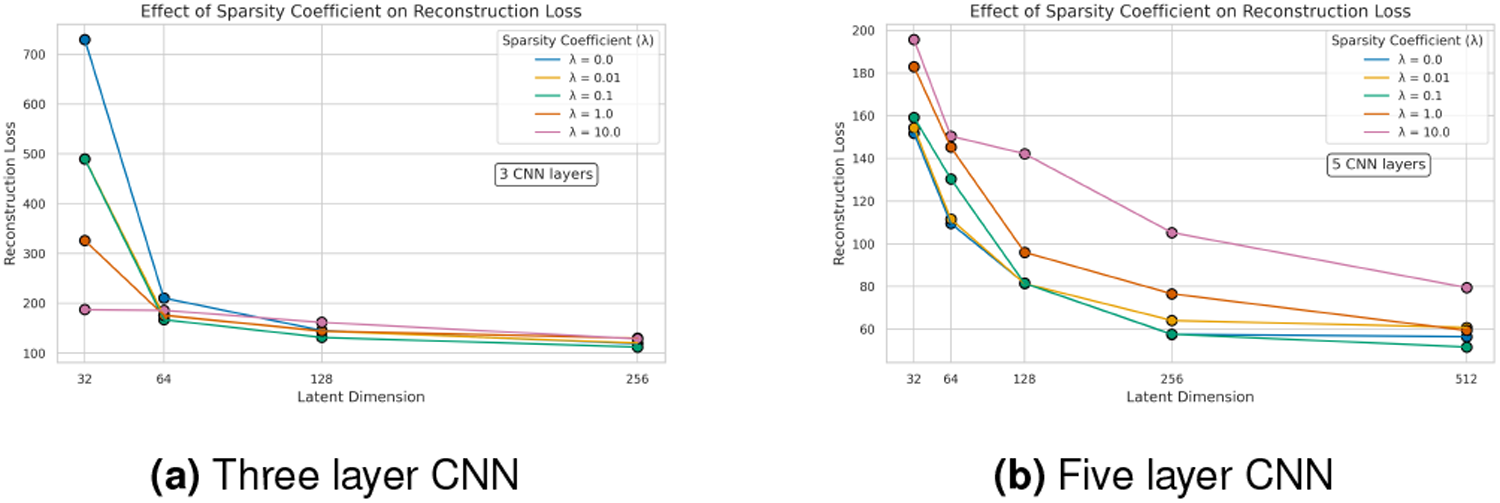
Effect of sparsity on reconstruction loss

### 4.2. Effect of hyperparameters on latent representation

To understand the impact of the hyperparameters on the quality of the generated embeddings, we used the synthetic dataset that contains 2D contours and corresponding labels (Sec. 3.1.3). We calculated the average Euclidean distances between the embeddings within a cluster. Embeddings from a cluster should be closer together. While this provides only an approximate measure of how different hyperparameters perform—since there are no “ideal” distances between embeddings in a cluster—it serves as a useful proxy for evaluating the performance of various hyperparameters.

For both CNN configurations, using sparsity loss greatly reduces the average distance between embeddings within a cluster (Fig. 4). This effect may be due to reduced activations of neurons in the latent space, enforced by the sparsity loss. The higher the sparsity coefficient, the lower the neuron activations, thereby leading to smaller Euclidean distances.

**Fig. 4.**
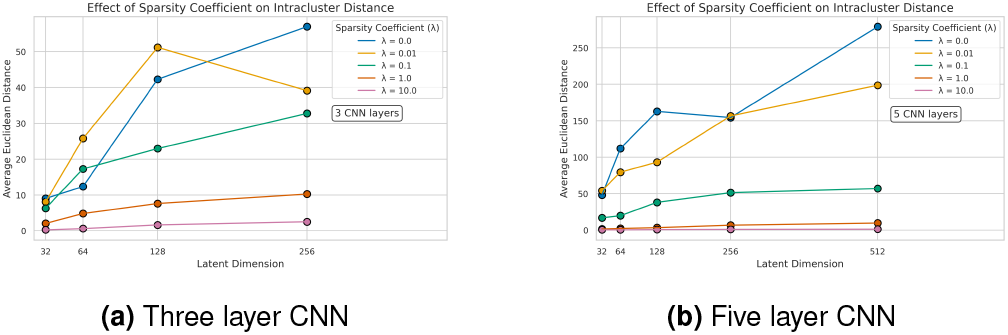
Effect of sparsity on intracluster distance

### 4.3. Cluster Analysis

For our clustering analysis, we selected the autoencoder model with the following configuration as it showed strong performance in the reconstruction loss on the test dataset (Sec. 4.1) and shorter average Euclidean distance on the synthetic dataset (Sec. 4.2):

- Number of CNN layers: 5
- Latent dimension: 512
- Sparsity coefficient: 1

Using the selected model, we first compare the performance of different clustering algorithms and then perform clustering analysis on TTG020-L dataset.

#### 4.3.1. Analysing performance of different algorithms

Using our synthetic dataset, we evaluated the performance of three clustering algorithms—DBSCAN, Agglomerative Clustering, and TAHC, using cosine similarity as the distance metric. We selected cosine similarity because it facilitates intuitive threshold tuning and is well-suited for comparing embedding vectors across different experiments.

For this analysis, we used the `scikit-learn`(29) implementations of DBSCAN and Agglomerative Clustering, and implemented TAHC as described in Section 3.3.1. For Agglomerative Clustering with cosine similarity, we explored two approaches: one using a predefined number of clusters, and another using a distance threshold. We did not include the adaptive mean-linkage algorithm (22) in our comparisons, as no official implementation is available. Additionally, their approach uses Euclidean distance, whereas our framework employs cosine similarity. We also found no evidence in the published work that the adaptive mean-linkage algorithm has been evaluated on high-dimensional datasets, which are characteristic of our experiments.

Clustering performance was assessed using the Adjusted Rand Index (ARI), a measure of similarity between predicted and true cluster assignments, adjusted for chance. Our proposed method, TAHC, correctly identified all clusters and outperformed the other algorithms (Table 1); hence, we used it for further analysis. In contrast, Agglomerative Clustering produced overlapping clusters—a limitation also observed with DBSCAN, even after moderate parameter adjustments. While iterative threshold adjustments in TAHC resolved these overlaps, Agglomerative Clustering failed to fully separate all clusters, and the number of detected clusters varied with minor threshold changes.

**Table 1.** Comparison of clustering algorithms on the synthetic dataset.

| Algorithm | # Clusters Defined | # Clusters Detected | Adjusted Rand Index (ARI) |
| --- | --- | --- | --- |
| DBSCAN | — | 13 | 0.74 |
| Agglomerative (fixed) | 16 | 16 | 0.66 |
| Agglomerative (threshold) | — | 21 | 0.82 |
| TAHC | — | 16 | **1.00** |

#### 4.3.2. Clustering gene expression profiles in TTG020-L

We performed clustering analysis on the TTG020-L gene expression dataset, which contains measurements from 45,101 probes. We filtered out probes whose maximum copy number (amplitude) was less than five, resulting in 17,055 gene expression profiles. We conducted the analysis both with and without incorporating the standard deviation. Using the method described in Sec. 3.4, we created four categories for variability. The following parameters for TAHC were selected based on the performance observed on the synthetic dataset, which served as a reference for subsequent clustering on the TTG020-L dataset.

**TAHC parameters**

- Cosine similarity threshold at first iteration: 0.85
- Threshold at succeeding iterations: 0.93, 0.94, 0.95, 0.97
- All subsequent iterations: 0.97
- Input: Embeddings

Using these parameters, we performed clustering on the TTG020-L dataset. To assess the performance of our framework and analyze the resulting clusters, we employed the Pearson correlation coefficient (PCC). Given the absence of labels and the lack of a predefined number of clusters, PCC was a suitable choice. Despite its limitations—such as sensitivity to outliers and the assumption of normal distributions—the Pearson correlation coefficient provides a useful approximation for understanding the clusters. Additionally, we used PCC as a point of comparison to evaluate the agreement between our clustering results and traditional correlation-based similarity. The generated clusters have an average PCC of around 0.8 (Fig.5) and include a mix of both larger and smaller clusters (Fig.6). Incorporating standard deviation resulted in an increase in the number of clusters, likely due to existing clusters being split based on the level of variability (Table 2).

**Table 2.** Comparison of PCC with and without incorporating variability.

| Method | Number of clusters formed | Mean PCC | Min PCC | Max PCC |
| --- | --- | --- | --- | --- |
| TAHC – without standard deviation | 1741 | 0.82 | 0.35 | 0.98 |
| TAHC – with standard deviation | 2289 | 0.81 | 0.28 | 0.99 |

**Fig. 5.**
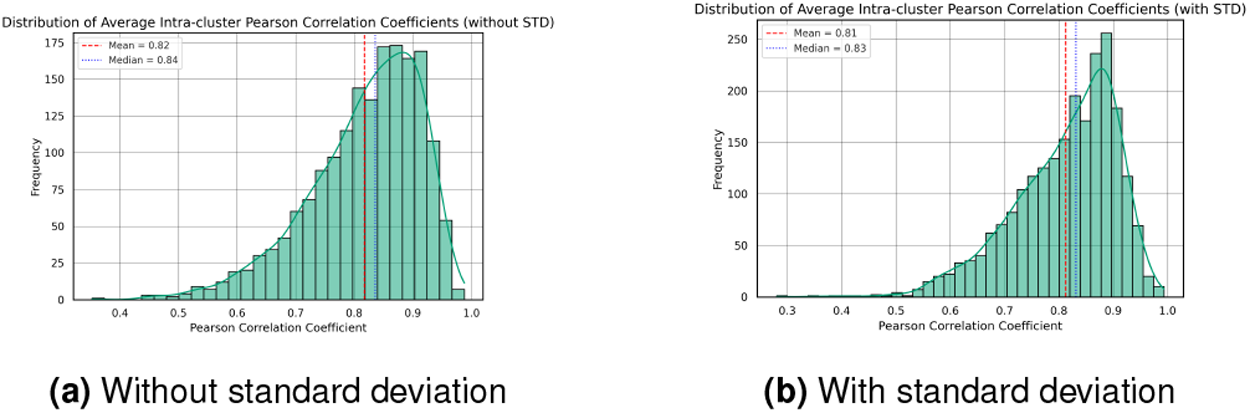
Histogram of average PCC

**Fig. 6.**
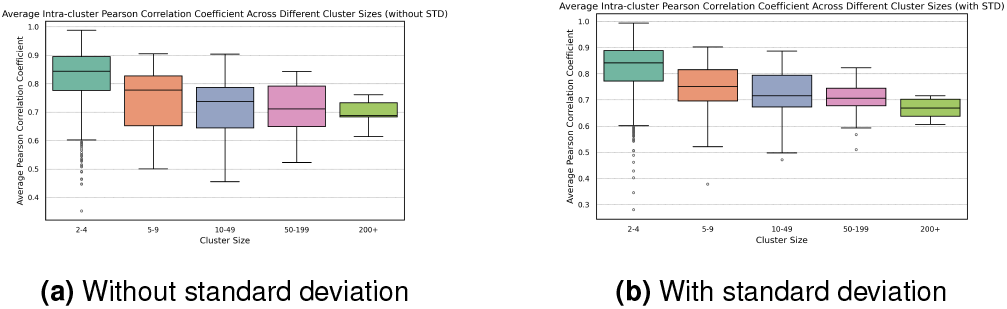
Boxplot of size of clusters

##### Visual analysis of select clusters

We next analyze several representative cases, comparing amplitudes, 2D contour plots, and 3D surface plots to evaluate how our approach performs against methods that rely solely on PCC. For this analysis, we used clustering methods applied to embeddings fused with standard deviation. We normalized amplitudes to the range [0, 1] to compare gene expression values for probes within the same clusters. We selected clusters with the highest and lowest PCC values; both were of size 2. Additionally, we chose a cluster with a PCC of approximately 0.83 and a size of 2. Although a PCC threshold of around 0.8 is commonly used, its selection lacks consensus, is often arbitrary, and may not reflect biological significance. (3).

##### Cluster with highest PCC (Fig.7)

**Fig. 7.**
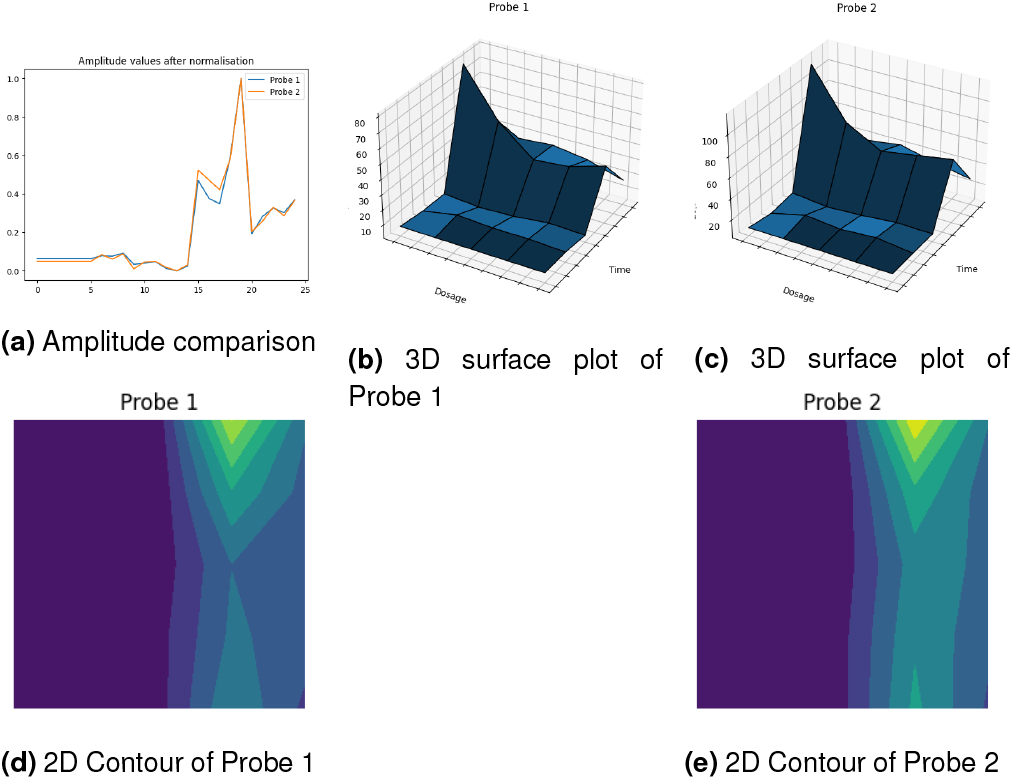
Cluster with highest PCC: 0.99

The cluster with the highest PCC (0.99349) contains two probes showing a matching trend. Both correspond to the gene symbol Cyp7a1, which encodes cholesterol 7 alpha-hydroxylase, a key enzyme in bile acid biosynthesis (30).

##### Cluster with PCC 0.83 (Fig.8)

**Fig. 8.**
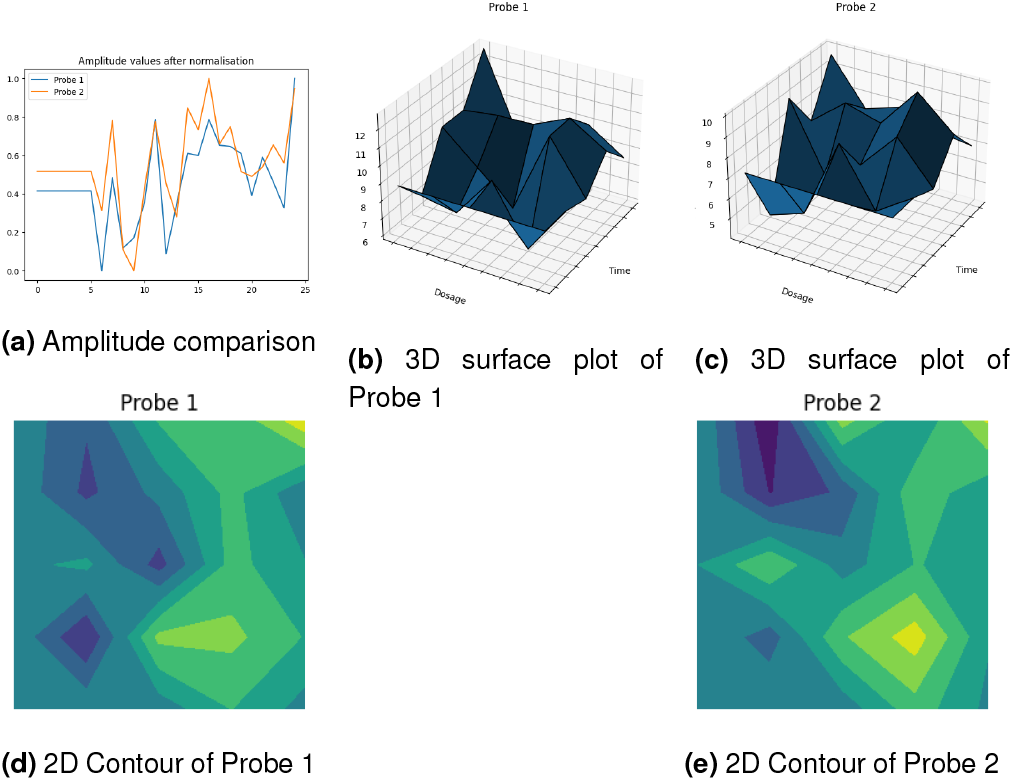
Cluster with PCC-0.83

In this case, most peak locations and the overall trends were well aligned between the two probes. Even after normalization, slight differences in amplitude magnitude may have contributed to the lower PCC; however, the general trend remained consistent across both profiles.

##### Cluster with lowest PCC (Fig.9)

**Fig. 9.**
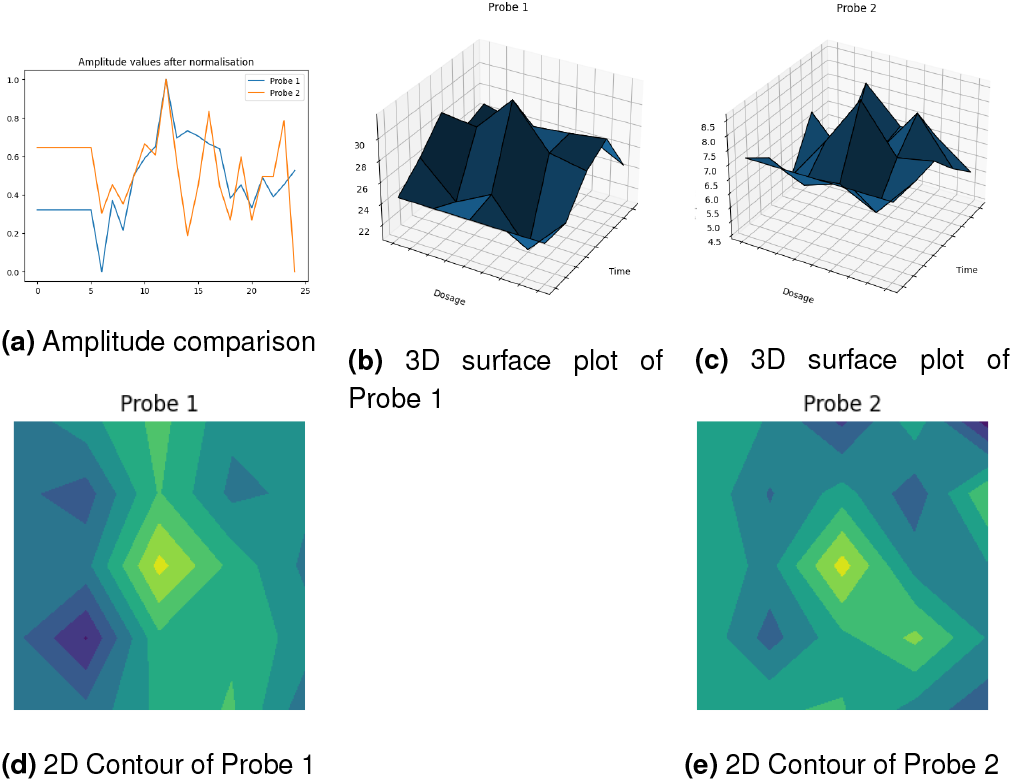
Cluster with lowest PCC 0.28

Despite the low PCC (0.28), the two gene expression profiles in this cluster exhibited a few peaks at the same positions, including the maximum peak.The probes correspond to TIMM10, a mitochondrial import chaperone, and TTC7B, a protein involved in phosphatidylinositol phosphate biosynthesis and plasma membrane protein localization.

#### 4.3.3. Functional Analysis of Gene Clusters Using KEGG Enrichment

To evaluate the biological relevance of the gene clusters derived from expression data, we performed functional enrichment analysis using the KEGG pathway database. This allowed us to identify whether specific gene clusters were significantly associated with known biological pathways. Out of 128 total clusters with more than 10 gene expression profiles, 32 (25.0%) showed enrichment for at least one KEGG pathway with an adjusted p-value less than 0.05.

A binary enrichment map was constructed to visualize the presence (p < 0.05) or absence of pathway enrichment across clusters (Fig. 10). This heatmap highlights clusters with extensive pathway-level coherence and shows that certain clusters harbor markedly more functional associations than others. To further quantify the degree of enrichment per cluster, we computed the number of significantly enriched pathways for each cluster with p < 0.05. The distribution is shown in Fig. 11a. While many clusters exhibited only a few enriched pathways, a small subset demonstrated substantial enrichment, suggesting a high level of internal functional organization. Among these, Cluster 1744 displayed a particularly strong enrichment profile, with 42 significantly enriched pathways. Fig.11b shows the distribution of adjusted p-values for all enriched pathways in this cluster, with a mean adjusted p-value of 0.02. This consistent enrichment signal suggests biological coherence rather than a statistical artifact. The average pairwise Pearson correlation coefficient (PCC) within the cluster was 0.68. Taken together, these results suggest that certain clusters—such as Cluster 1744—are functionally closed-form, meaning they exhibit a high degree of internal coherence both statistically and biologically. These clusters may represent meaningful transcriptional modules that align closely with canonical pathways and could be prioritized for further mechanistic investigation.

**Fig. 10.**
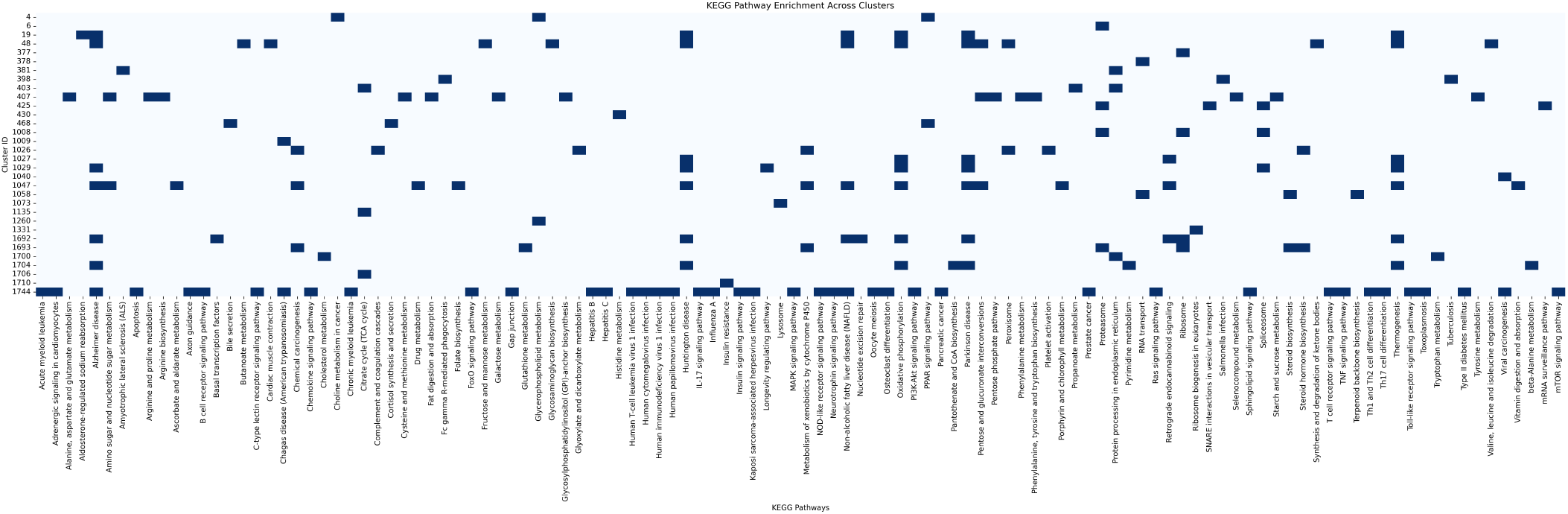
Binary enrichment map showing presence (*p <* 0.05) or absence of KEGG pathway enrichment across clusters. Each row corresponds to a cluster and each column to a KEGG pathway.

**Fig. 11.**
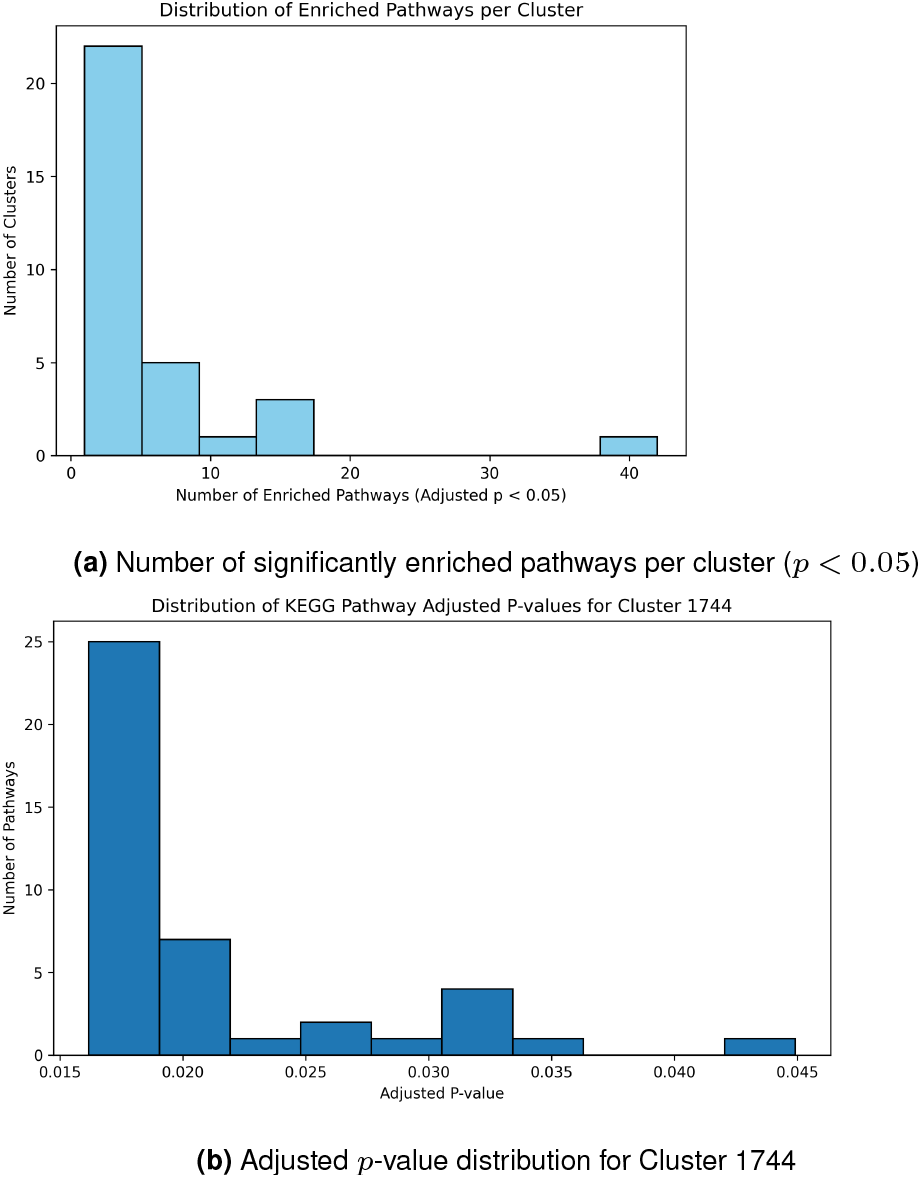
Functional enrichment p-value distributions

## 5. Conclusion

In this paper, we presented a new methodology that incorporates the visual features of gene expression plots and accounts for variability to identify co-expressed genes in the Percellome database. While our approach was specifically tested on the Percellome project, it is generalizable and can be applied to other datasets containing gene expression profiles that vary across two experimental conditions, such as dosage and time.

## Acknowledgment

This work was supported by Comprehensive Study on PFASs organized by the Ministry of the Environment, Japan.

